## Supplementary Figure 1 for "OpenNeuro: An open resource for sharing of neuroimaging data"

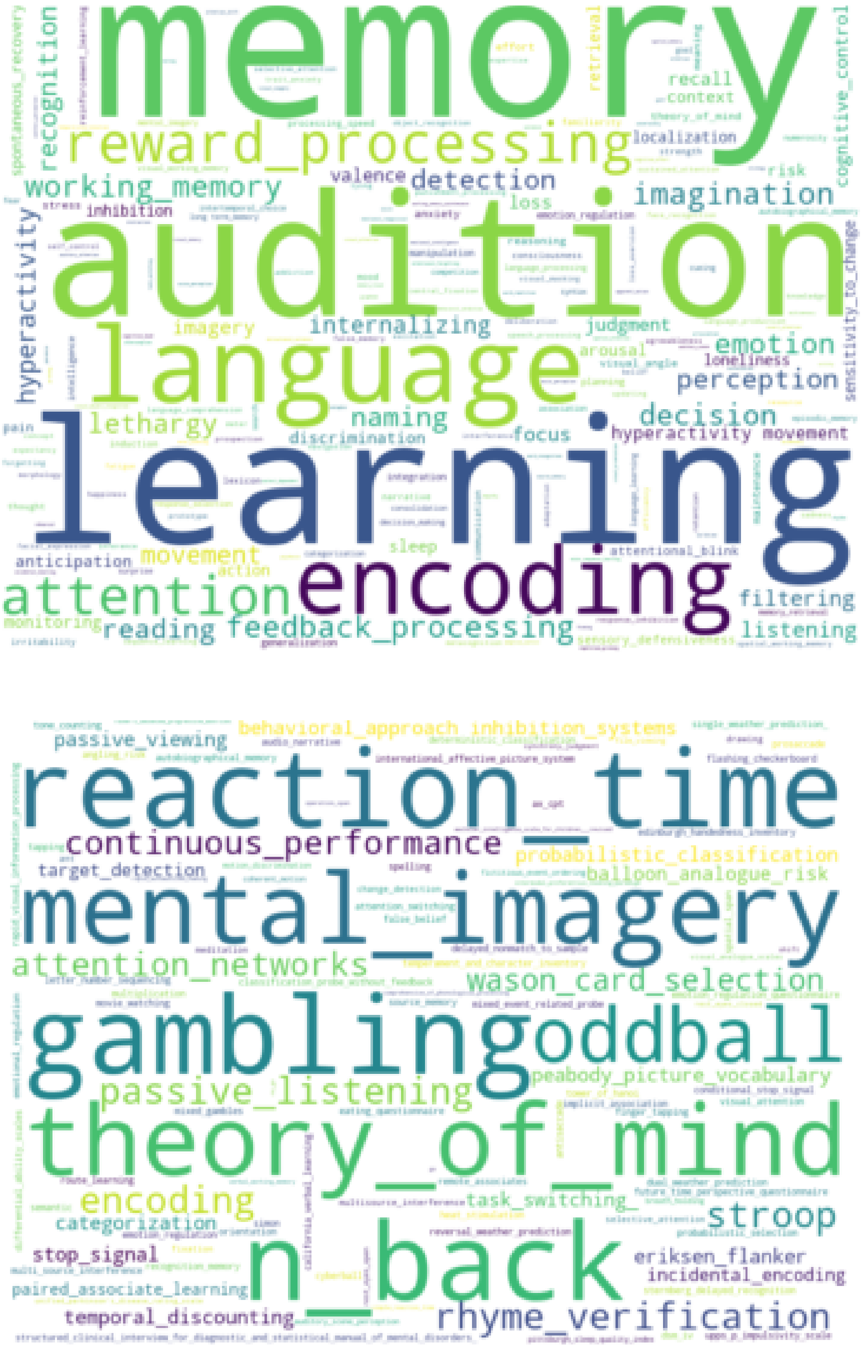


*Supplementary Figure 1*. Word clouds based on Cognitive Atlas terms for psychological concepts (top) and tasks (bottom) identified from titles and README files associated with OpenNeuro datasets.
